# The gastric mucosa of Atlantic salmon (*Salmo salar*) is abundant in highly active chitinases

**DOI:** 10.1101/2022.05.10.491315

**Authors:** Matilde Mengkrog Holen, Tina Rise Tuveng, Matthew Peter Kent, Gustav Vaaje-Kolstad

**Affiliations:** Center for Integrative Genetics, Department of Animal and Aquacultural Sciences, Faculty of Biosciences, Norwegian University of Life Sciences, Ås, Norway; Faculty of Chemistry, Biotechnology and Food Science, Norwegian University of Life Sciences, Ås, Norway

**Keywords:** chitin, chitinase, gastric mucosa, Atlantic salmon

## Abstract

The Atlantic salmon (*Salmo salar*) genome contains 10 chitinase encoding genes, but little is known about the function of these chitinases. We show that the protein products of three genes, the family 18 glycoside hydrolase (GH18) chitinases Chia.3, Chia.4, and Chia.7 are secreted in the stomach mucosa and are amongst the most abundant proteins in this matrix. Chia.3 and Chia.4, sharing 95% sequence identity, were not possible to separate by standard chromatographic methods and were thus purified as a chitinase pair. Biochemical analysis revealed chitinolytic activity towards β-chitin for up to 24 hours at pH 2-6. Further *in vitro* analysis showed that this chitinase pair efficiently degraded various chitin-containing substrates to chitobiose (GlcNAc_2_) suggesting that Atlantic salmon has the potential to utilize novel chitin-containing feed sources.

## INTRODUCTION

Chitin is an insoluble polysaccharide, consisting of β-1,4-linked *N*-acetyl-D-glucosamine residues. It is one of the most common biopolymers in nature and is found in three distinct allomorphic forms depending on the orientation of the chitin chains ^1^: α-chitin, β-chitin, and γ-chitin. In α-chitin, the most abundant form of chitin, the polymer chains have antiparallel orientation resulting in stronger intermolecular forces compared to β-chitin and γ-chitin with a parallel and a mixture of parallel and antiparallel orientation of the polymer chains respectively. Chitin functions as a structural component in algae and fungi cell walls ^2,3^, in the exoskeleton of arthropods such as crustaceans and insects ^4,5^, and is even hypothesized to be present in the scales and gut lining of some vertebrates, including ray-finned fish ^6,7^.

Chitinases are thought to play a role in at least three different processes ^8^, firstly in the breakdown of chitinous body structures during development, secondly they can be deployed to defend against infection of chitinous pathogens, and thirdly they can be involved in the digestion of chitin for nutrient absorption and energy production. Fish are known to express chitinases from the glycoside hydrolase 18 family (GH18), a multigene family with a conserved catalytic motif; DXXDXDXE ^9^, but little is known about the role of these chitinases. To our knowledge, Jeuniaux (1961) was the first to report endogenous chitinase activity in the fish gut using a β-chitin suspension from squid pen as substrate ^10^. Such gut chitinases are hypothesized to be secreted by the gastric mucosa ^11^. These enzymes have been shown to have an acidic pH optimum, while fish that lacks an acidic stomach (e.g. zebrafish) have comparable activities at neutral pH ^12^. Chitinase activity in fish intestines has not been shown to correlate with the amount of dietary chitin, but fish that swallow prey whole have shown higher chitinase activity relative to the fish that have pharyngeal teeth or other buccal cavity modifications ^13^. Furthermore, fish chitinases are mainly expressed in the stomach, and stomach chitinases have different activities on various insoluble chitin substrates depending on diet ^14^. This indicates that fish chitinases could aid in the digestion of chitinous feed.

Wild Atlantic salmon (*Salmo salar*) is known to prey on chitinous organisms such amphipods, euphausiids, shrimp, and insects ^15,16^ and possess 10 genes encoding family GH18 chitinases ^17^. A better understanding of the biological functions of these proteins may be of value to the salmon industry as it searches for new, alternative feed sources. Here, we quantify the relative amount of stomach chitinases in the gastric mucosa of Atlantic salmon and isolate and characterize two of these. The results provide evidence that chitin can be degraded by these enzymes within the salmon gut.

## RESULTS

### Sequence analysis

The Atlantic salmon genome encodes 10 chitinase-like genes according to the NCBI RefSeq annotation (GCF_000233375.1; release 100) that all belong to family 18 of the glycoside hydrolases, as classified by the carbohydrate active enzyme (CAZy) database ^18^. Three of these chitinase genes (hereby named *chia.3, chia.4*, and *chia.7*; proteins named Chia.3, Chia.4, and Chia.7, respectively) have shown stomach-specific expression ^17^. The sequence identity of the amino acid sequences of the chitinases range from 60-95% when aligned with each other and 61-65% when aligned with the ortholog human acidic mammalian chitinase, AMCase (Table 1).

**Table 1.** Sequence identity (%) between Chia.3, Chia.4, and Chia.7 and AMCase (the UniProt ID is given in parenthesis) after removing the predicted signal peptide sequence (identified with SignalP v.5.0 ^19^)

|  | **Chia.3**<br>**(A0A1S3L8D8)** | **Chia.4**<br>**(A0A1S3L8T9)** | **Chia.7**<br>**(A0A1S3MFN1)** | **AMCase**<br>**(Q9BZP6)** |
| --- | --- | --- | --- | --- |
| **Chia.3**<br>**(A0A1S3L8D8)** | 100 | 95 | 60 | 65 |
| **Chia.4**<br>**(A0A1S3L8T9)** | 95 | 100 | 60 | 65 |
| **Chia.7**<br>**(A0A1S3MFN1)** | 60 | 60 | 100 | 61 |

All three chitinases share an N-terminal signal peptide, the GH18 catalytic motif DGLDXDWE, multiple putative *N*-Acetylgalactosamine (GalNAc) O-glycosylation sites, and a C-terminal domain identified as a family 14 carbohydrate-binding module (CBM14; identified by dbCAN2 annotation ^20^) (Figure 1). In addition, AMCase, Chia.3 and Chia.4 share three residues hypothesized to be important for acidic activity ^21,22^: an arginine (R) at position 145 and histidine (H) residues at positions 208 and 269. Chia.7 shows the least sequence identity to AMCase and has asparagine (N) residues at position 208 and 269.

**Figure 1.**
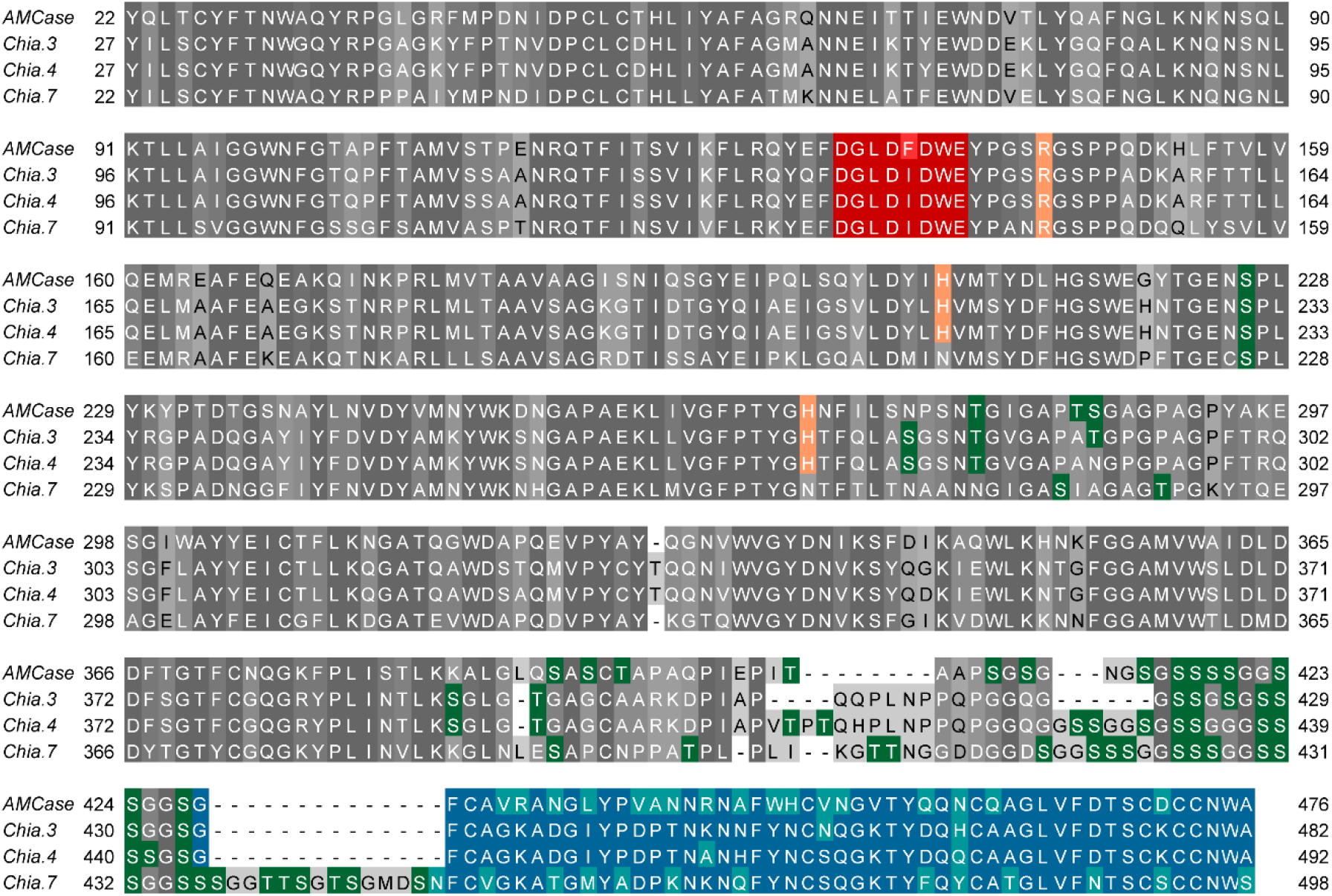
Multiple sequence alignment of stomach-specific chitinases from Atlantic salmon (Chia.3, Chia.4 and Chia.7) with human acidic mammalian chitinase (AMCase). The N-terminal signal peptides (identified with SignalP v.5.0 ^19^, default settings) were removed from the alignment. The catalytic motif residues are highlighted in red. Residues in the catalytic domain that may be important for activity in acidic environments are highlighted in orange. Predicted GalNAc O-glycosylation sites (identified with NetOGlyc v.4.0 ^23^, default settings) are highlighted in green and the C-terminal CBM14 (identified with dbCAN2 annotation ^20^) residues are highlighted in blue (cyan if the sequence identity is low).

### Chitinase abundance correlate with gene expression levels

Label-free quantitative (LFQ) proteomics was used to determine the relative amount of proteins in the gastric mucosa of Atlantic salmon collected after one day without feed. Strikingly, Chia.3, Chia.4, and Chia.7 were among the most abundant proteins in the gastric mucosa (Figure 2; Supplementary Table 1). We took advantage of published RNA-seq data generated from Atlantic salmon stomach tissue (ArrayExpress, E-MTAB-10594) to determine the correlation between the relative amount of the top 20 secreted stomach proteins and the gene expression levels of the genes coding for these proteins. The Spearman correlation (ρ) between the relative amount of proteins (log_2_(LFQ intensity + 1)) and gene expression levels (log_2_(TPM + 1), where TPM stands for transcripts per million) was 0.68 (p-value = 0.0014), indicating a positive correlation between gene expression levels and protein abundance of the secreted stomach proteins (Figure 2). Eight genes were both the most highly expressed transcripts and the most abundant proteins in the stomach including the three chitinases (Chia.3, Chia.4, and Chia.7), two Pepsin-A-like proteases, one IgGFc-binding protein-like, a cysteine protease inhibitor; cystatin, and a lectin; fish-egg lectin. The full proteomic dataset is available in Supplementary Table 2.

**Figure 2.**
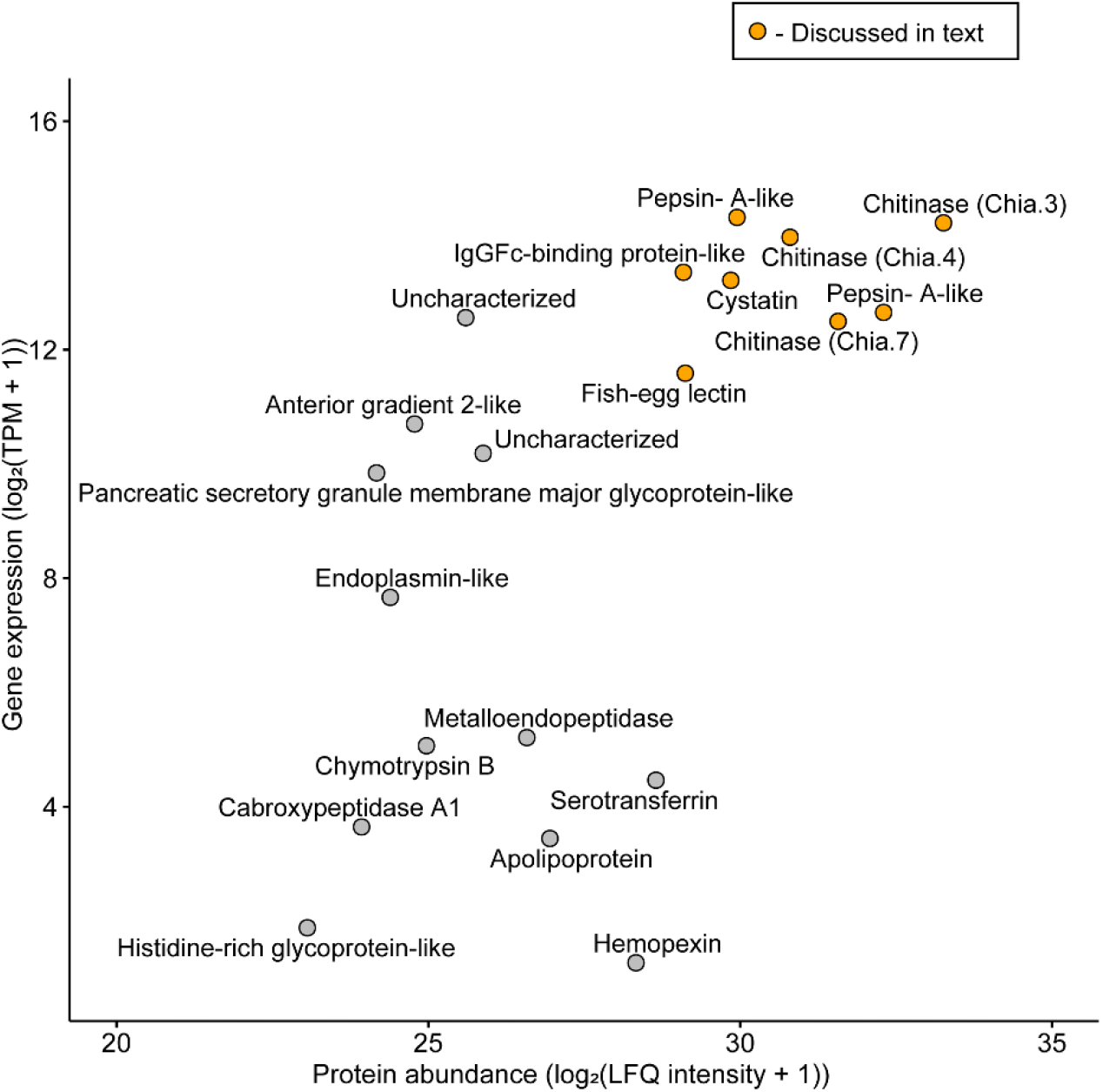
Spearman correlation between protein abundance of proteins (mean of log_2_(LFQ intensity + 1), n = 3) and gene expression levels (mean of log_2_(TPM + 1), n = 13) in Atlantic salmon stomach. The points highlighted in orange are discussed in the text. The corresponding Uniprot IDs can be found in Supplementary Table 1.

### Purification and characterization of Atlantic salmon stomach chitinases

To capture the Atlantic salmon stomach chitinases, a stomach tissue homogenate was passed over a chitin-affinity column and the bound proteins were eluted by pH reduction from 7 (binding buffer) to 3 (elution buffer). The eluate contained two proteins represented by two distinct bands at approximately 50 and 45 kDa (Figure 3). The identity of the proteins was determined by LC-MS/MS, showing the presence of Chia.3 and Chia.4 in both bands. The predicted molecular weights of Chia.3 and Chia.4 after removal of the N-terminal signal peptide are 49.6 and 50.4 kDa respectively, and the 5 kDa difference of the two bands corresponds to the molecular mass of the CBM14 domain. The proteins were not possible to separate by standard chromatographic techniques, most likely due to their highly similar protein sequence (95% alike).

**Figure 3.**
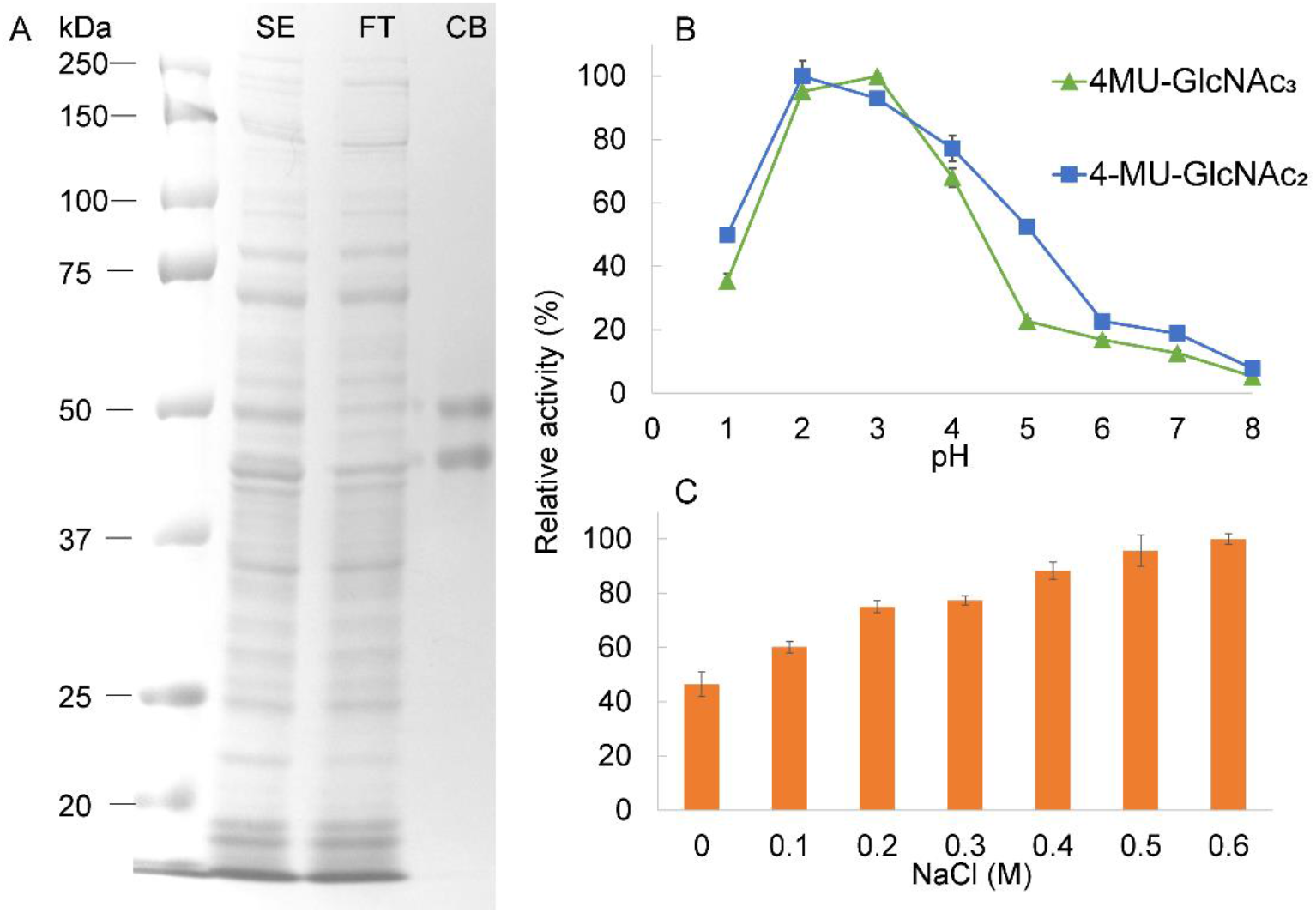
Purification and characterization of stomach chitinases from Atlantic salmon. (**A**) SDS-PAGE analysis of Atlantic salmon stomach chitinases were purified using chitin affinity chromatography; (1) ladder (Precicion Plus Protein, BioRad), (2) SE: total stomach extract, (3) FT: column flow-through, (4) CB: proteins eluted from the washed chitin affinity column. (**B**) Enzymatic activity at different pH values, using 100 μM 4MU-GlcNAc_2_ (blue) and 100 μM 4MU-GlcNAc_3_ (green) as substrate, with enzyme concentrations of 31 and 7.8 nM respectively. The reactions were incubated for 15 minutes at 37 °C in 0.1 M Gly-HCL buffer (pH 1.0 – 3.0) or McIlvaine’s buffer (0.1 M citric acid and 0.2 M Na_2_HPO_4_, pH 4.0 – 8.0). Data points are joined by lines to indicate the shape of the pH activity relationship (**C**) Enzymatic activity with increasing NaCl concentration (0-0.6 M) using 100 μM 4MU-GlcNAc_2_ as substrate and an enzyme concentration of 31 nM, with conditions identical as in (B), incubating the reaction in McIlvaine’s buffer at pH 5. The activity was assessed in triplicate, values shown are means ± SD.

Chia.7 was not detected in the protein eluted from the column, but the fact that we know the protein is abundantly expressed suggests that conditions used in this affinity purification are not suited to capture the Chia.7 protein. All subsequent enzyme assays were performed on the Chia.3 + 4 eluate.

To determine the influence of pH and NaCl on the activity of the chitinases, activity was determined using the chitotriose analog 4MU-chitobioside (4MU-GlcNAc_2_) or chitotetraose analog 4MU-chitotrioside (4MU-GlcNAc_3_). The highest activity was observed at pH 2 and 3 followed by a gradual decrease to pH 8 where essentially no activity could be measured (Figure 3B). The addition of NaCl to the reaction mixture yielded a chitinase activity that increased with increasing salt concentrations, showing almost a doubling in activity from 0 to 0.6 M NaCl (Figure 3C). The latter salt concentration approximately represents the salinity of seawater.

### Activity towards insoluble chitin

Information about the pH optimum of an enzyme is sometimes not predictive of the enzyme performance over time (operational stability). To determine the hydrolytic potential of the chitinase pair in relevant pH conditions using a relevant substrate, the progress of a chitin degradation reaction was followed for 24 h in reactions having pH ranging from 2-6 (Figure 4). Interestingly, the highest yield after 24 h was obtained at pH 5, for which the pH optimum was less than 50% for that of pH 2 and 3 (Figure 3, panel B). The major product arising from the reactions was chitobiose (GlcNAc_2_), but some GlcNAc was also observed (<10% of the total products).

**Figure 4.**
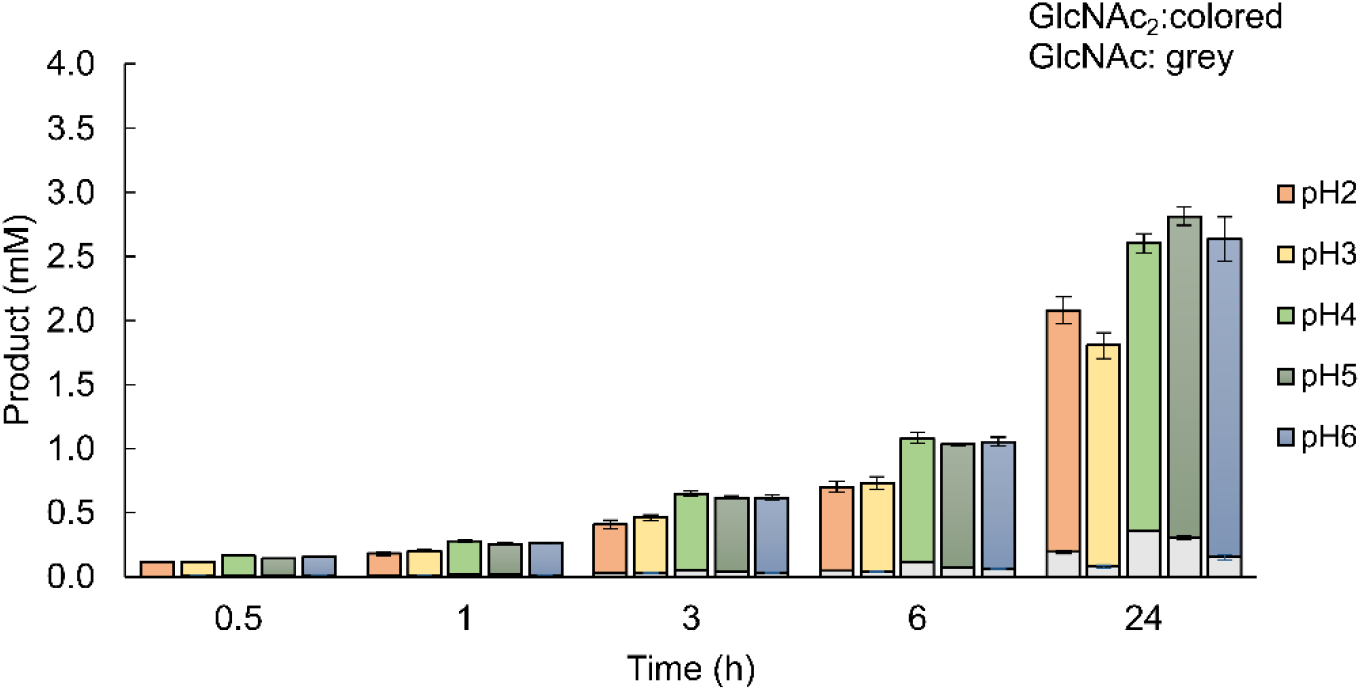
Operational pH stability. Concentrations of degradations products, GlcNAc dimers (GlcNAc_2_, colored boxes) and monomers (GlcNAc, grey boxes), produced by Chia.3 + 4 when incubated with β-chitin in pH 2-6 over 24 hours are shown in a bar chart representation. The amount of degradation products was analyzed after 0.5, 1, 3, 6, and 24 hours of incubation at 14 °C. The reactions were performed with an enzyme concentration of 0.2 μM and a substrate concentration of 10 mg/mL. All reactions were run in triplicates, values are means ± SD.

### The activity of Atlantic salmon stomach chitinases and ChiB from Serratia Marcescens on α-chitin-containing organisms

The ability of the chitinase pair to depolymerize insoluble β-chitin prompted us to investigate whether the enzymes were able to break down α-chitin-containing organisms commonly found in the diet of Atlantic salmon using the shell from shrimp and crab, and skin from black soldier fly pupae. To put the activity of the chitinase pair in a metabolic context, the well-characterized chitinase from the soil bacterium *Serratia marcescens, Sm*ChiB was included for comparison. The experiment was done at 14 °C and pH 4.8 which are conditions similar to the Atlantic salmon stomach environment ^24^.

The results show that the chitinase pair and *Sm*ChiB were all able to partly depolymerize the chitin-containing substrates tested (Figure 5). The Atlantic salmon chitinase pair showed substantial activity to all substrates investigated except the dried shrimp shell and showed higher chitinolytic activity on all substrates compared to *Sm*ChiB under Atlantic salmon gastric-like conditions, i.e. not optimal conditions for *Sm*ChiB ^25^. Interestingly, *Sm*ChiB was less active on purified α-chitin than crab shell, while the opposite was observed for the chitinase pair after 6 hours of incubation.

**Figure 5.**
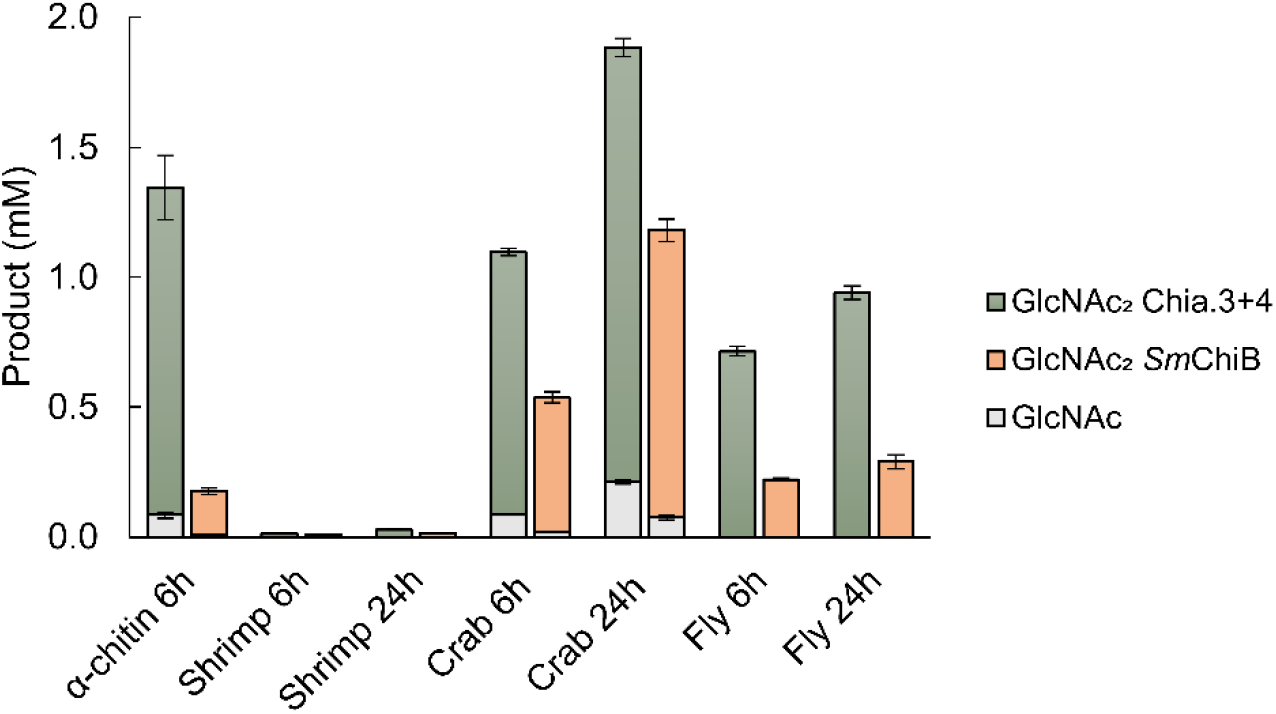
Degradation of natural α-chitin substrates. The bar chart displays the amount of degradation products, GlcNAc dimers (GlcNAc_2_, colored boxes) and monomers (GlcNAc, grey boxes), produced by Atlantic salmon stomach chitinases (Chia.3 and Chia.4) compared to ChiB from *Serratia marcescens* (*Sm*ChiB) when incubated with α-chitin for 6 hours and shrimp shell (shrimp), crab shell (crab) and black soldier fly pupae skin (fly) for 6 and 24 hours in 0.1 M sodium acetate buffer (pH 4.8) at 14 °C. The reactions were performed with an enzyme concentration of 0.5 μM and a substrate concentration of 10 mg/mL for all substrates except black soldier fly pupae skin with a substrate concentration of 25 mg/mL. All reactions were run in triplicates, values are means ± SD.

## DISCUSSION

Atlantic salmon possess multiple genes encoding chitinases including three variants that are abundant as both gene transcripts and proteins in the stomach. The active proteins are secreted into the gastric mucosa but their functional role there is unknown. The stomach of Atlantic salmon works both as a “feed-grinder” and as a first-line defense against water- and feed-borne pathogens. In theory, chitinases could play a role in the digestion of chitin-containing feed and/or chitin-containing pathogens such as fungi. The results show that the chitinases are present together with high levels of IgGFc-binding-like protein, cystatin and fish-egg lectin (Figure 2), three proteins shown to play a potential role in the teleost immune system against pathogens ^26–30^. Moreover, the Atlantic salmon stomach chitinases share many features with human AMCase ^31^; they share the same conserved domains, they are abundant in the gastrointestinal tract, they are acid-stable, and they are able to degrade crab shell chitin. AMCase has shown to play potential roles in digestion and the immune response against chitin-containing organisms, a role that could be similar for the Atlantic salmon gastric chitinases.

In this study, Chia.3 and Chia.4 were purified from Atlantic salmon stomachs using chitin affinity chromatography, but we were unable to purify Chia.7 despite its identification by proteomic analysis. Chia.3 and Chia.4 are almost identical, while Chia.7 shows less sequence identity to the other stomach chitinases and differences in chitin-binding capacities between the different chitinases could be the reason for not being able to isolate Chia.7 in the same conditions. The combined fraction of pure Chia.3 and Chia.4 showed a pH optimum of pH 2 and 3 respectively when using fluorogenic substrates (4MU-chitobioside and 4MU-chitotriose). This pH optimum was different from the operational pH stability optimum of pH 5 determined towards β-chitin, a more realistic substrate for the chitinases. Differences in substrate length and composition have previously been shown to affect the chitinolytic pH optimum ^32^, and this underscores the need to carefully consider the substrates used when characterizing proteins. Also, the non-natural aglycon leaving group of the 4MU-conjugated substrates and the insoluble nature of β-chitin may contribute to the pH preference of the enzyme. Atlantic salmon has a gastric stomach with an average pH of 4.8 depending on both feed type and time since ingestion ^24^, which corroborates the operational pH stability data. Further, the pH in the contents of the most distal part of the stomach is generally higher (pH 5) than the middle part and increases to a more neutral pH (pH 8) in the pyloric, mid, and distal intestine. Our results show that the Atlantic salmon stomach chitinase pair were highly active and stable at pH 2.0-6.0 over 24 hours. This is biologically relevant as it may take up to somewhere between 24 to 48 hours for the fish to empty its stomach upon ingestion ^33^. A pH optimum of pH 2 when using fluorogenic substrates has previously been reported for stomach chitinases isolated from humans, mice, chicken, and pigs ^31,34–36^. Moreover, the optimum pH of stomach chitinases from Silver croaker (*Pennahia argentatus*), Marbled rockfish (*Sebastiscus marmoratus*), Red sea bream (*Pagrus major*), Japanese eel (*Anguilla japonica*), and Red scorpionfish (*Scorpaena scrofa*) have been reported to be pH 4.0-5.0, 4.0-4.5, 5.5, 4.4, and 5.0 respectively when using colloidal chitin as a substrate ^14^. This is in line with our observations.

Salmon migrate from the rivers into the sea and must adapt to a change in diet and increasing salt concentrations. The results show that the stomach chitinases of Atlantic salmon were substantially more active at 0.6 M NaCl, equivalent to seawater salinity, than without any NaCl. The NaCl concentration of seawater corresponds to an osmolality of 1200 mOsm and the stomach chyme osmolality of rainbow trout (*Oncorhynchus mykiss*) has been reported to reach a maximum of 775 mOsm two hours after feeding ^37^. This value was reported for fish living in freshwater, and it is probably higher in seawater salmonids since it is known that salmon drink more external water for osmoregulation in seawater than in freshwater and because the gut is an important osmoregulatory organ in seawater teleost ^38,39^. Furthermore, our results show that the Atlantic salmon stomach chitinases degraded chitin from shrimp shells, crab shells and black soldier fly pupae more efficiently than ChiB from *S. marcescens*. The latter bacterium utilizes chitin as an energy source and is one of the most efficient chitin degraders out of 100 tested microorganisms ^40,41^. The higher efficiency of salmon chitinases could be a result of non-optimal conditions for *Sm*ChiB which works best at pH 5.0-6.0 with a temperature optimum of 58 °C ^25^, and/or a synergy effect of the two salmon chitinases working together.

Recent studies show that insect meal from black soldier fly has the potential to replace fish meal in the aquaculture industry ^42,43^ and that chitin and chitin degradation products can act as immunostimulants ^44,45^. Altogether, our results show that using chitin-containing organisms as novel feed sources for farmed Atlantic salmon can be of nutritional value.

## CONCLUSION

Our results show that some of the most dominant proteins in the stomach of Atlantic salmon are chitinases that are capable of effectively degrading chitin or chitin-containing substrates from various sources. The stomach chitinases are active and stable in the gastric-like conditions of Atlantic salmon and are therefore likely to play a role in the digestion of chitin-containing organisms commonly found in the natural diet of salmon. The results presented here can be taken into consideration when searching for novel feed ingredients in the aquaculture industry.

## ACKNOWLEDGMENTS

We want to thank Morten Skaugen at the Proteomics Core Facility at NMBU for helping with sample preparation and mass spectrometry analysis. We are also grateful to the people at the Center for Sustainable Aquaculture at NMBU for providing us with the fish used in the experiments and Sophanit Mekasha for providing us with *Sm*ChiB.

## METHODS

### Multiple sequence alignment

The amino acid sequences of Chia.3, Chia.4, Chia.7, and AMCase were downloaded from Uniprot and the alignment was calculated using Mafft with default settings in Jalview v. 2.11.14. The signal peptides were predicted with SignalP v.5.0 ^19^ using default settings. The GalNAc O-glycosylation sites were identified with NetOGlyc v.4.0 ^23^ using default settings and the CBM14 domain was identified with dbCAN2 annotation ^20^.

### Chitinous substrates

β-chitin (extracted from squid pen, Batch 20140101, France Chitin, Orange, France) was pulverized with a bead mill (Planetary Ball Mill PM 100, Retsch) to approx. 200 μm particle size. Shrimp shell was peeled off shrimps (*Pandalus Borealis*, Polar Seafood Norway) and the filling was removed from crab shell (*Cancer pagurus*, Lofotprodukt). Both products were bought at the local food market. Shrimp shell, crab shell, and black soldier fly pupae skin (*Hermetica illucens*, a kind gift from Fraunhofer-Gesellschaft, Munich, Germany) were dried at 105 °C overnight before the experiments were run. Shrimp- and crab shells were first crushed with mortar and pestle before the shells, black soldier fly pupae skin, and α-chitin (extracted from *Pandalus borealis*, Seagarden, Avaldsnes, Norway) were pulverized with a bead mill (Planetary Ball Mill PM 100, Retsch) to approx. 200 μm particle size.

### Enzymes

Chitinases from Atlantic salmon were isolated from stomach tissue and purified as described below. Chitinase B from *Serratia marcescens* (*Sm*ChiB) was overexpressed in *Escherichia coli* and purified as previously described ^46^.

### Proteomic analysis of Atlantic salmon gastric mucosa

Gastric mucosa of adult Atlantic salmon (n = 3, two male and one female, average fish weight 2245 g) was obtained from the process plant for fish farming laboratory at The Norwegian University of Life Sciences (NMBU). The fish was starved for one day before they were euthanized by a blow to the head. The stomach was dissected from the fish, the gastric mucosa was scraped off and mixed with 1 mL ice-cold phosphate buffer (20 mM sodium phosphate buffer pH 7.0, 150 mM NaCl) with 1X protease inhibitor cocktail (complete, EDTA-free Protease Inhibitor Cocktail, Roche) by pipetting and gentle vortexing. The homogenate was centrifuged at 12,000 rpm for 10 minutes at 4 °C and the supernatant was filtered through a 40- μm cell strainer. The total protein concentration was determined with Bradford Protein Assay (Bio-Rad) using Bovine Serum Albumin as standard. A total amount of 2 μg protein was loaded on an SDS-PAGE gel. The proteins were allowed to enter the gel, but without full separation of the proteins in the gel. The gel was stained with Coomassie Brilliant Blue R-250 (Bio-Rad) and de-stained before the region of the gel containing proteins were cut out for in-gel digestion, essentially performed as described by Shevchenko et al ^47^. In brief, proteins were reduced with DTT and alkylated with iodoacetamide before trypsinization. ZipTip C_18_ pipette tips (Merck Millipore) were used to purify peptides, followed by drying under vacuum. The peptides were dissolved in 10 μl of 2% (v/v) acetonitrile, 0.1% (v/v) trifluoroacetic acid and the peptide concentration was measured using av NanoDrop One and used to normalize the amount of peptides injected for LC-MS/MS analysis. The LC-MS/MS analysis was performed as described by Tuveng et al ^48^.

MS Raw files were analyzed using MaxQuant ^49^ version 1.6.17.0, and proteins were identified and quantified using the MaxLFQ algorithm ^50^. Samples were searched against the proteome of *Salmo salar* downloaded from Uniprot (UP000087266), and a list of common contaminants (included in the MaxQuant software package). As variable modifications protein N-terminal acetylation, oxidation of methionine, conversion of glutamine to pyroglutamic acid, and deamination of asparagine and glutamine were used, while carbamidomethylation of cysteine residues was used as a fixed modification. Trypsin was used as a digestion enzyme and two missed cleavages were allowed. The feature ‘Match between runs’ in MaxQuant, which enables identification transfer between samples based on accurate mass and retention time, was applied with default settings ^50^. The results from MaxQuant were further processed using Perseus (version 1.6.15.0) Proteins categorized as ‘only identified by site’, ‘reverse’ or as ‘contaminant’ were removed from the dataset. As an additional cut-off criterium, proteins were only considered present if they were detected in at least two of three replicates. The LFQ (label-free quantification) intensities were log_2_-transformed and averaged before analysis. The downstream analysis focused on proteins predicted to have a signal peptide using the *Salmo Salar* proteome (UniProt id: UP000087266).

### Comparison of protein abundance with gene expression levels

RNA-sequencing data from the stomach of Atlantic salmon was downloaded from ArrayExpress under project number E-MTAB-10594. The bcbio-nextgen pipeline (https://github.com/bcbio/bcbio-nextgen was used to trim, map and count raw reads before aligning to the Atlantic salmon genome (ICSASG_v2) ^51^ using STAR ^52^. Reads aligned to genes were counted with FeatureCounts ^53^ and transformed to transcripts per million (TPM) values to normalize for gene length. The TPM values were log_2_-transformed and averaged (n = 13) before analysis. A subset of genes coding for the 20 most abundant proteins secreted in stomach mucosa was used to calculate the Spearman correlation between the gene expression levels (log_2_(TPM + 1)) and the relative protein abundance (log_2_(LFQ intensity + 1)). The statistical analysis was done using the “ggpubr” package in R v.4.0.3.

### Purification of chitinases from stomach tissue

Stomach tissue (n = 2, average 7.2 g per purification, two rounds of purification) from adult Atlantic salmon (one female, one male, average weight 2230 g) was obtained from the process plant for the fish farming laboratory at NMBU. The fish was euthanized by a blow to the head and the stomach was dissected from the fish. Stomach content was removed before the tissue was cut into small pieces and stored in ice-cold phosphate buffer (20 mM sodium phosphate buffer pH 7.0, 150 mM NaCl) with 2X protease inhibitor cocktail (complete, EDTA-free Protease Inhibitor Cocktail, Roche). The tissue was homogenized directly after dissection.

Stomach tissue was homogenized in 5 volumes of ice-cold phosphate buffer (20 mM sodium phosphate buffer pH 7.0, 150 mM NaCl) with 2X protease inhibitor cocktail (complete, EDTA-free Protease Inhibitor Cocktail, Roche) using TissueRuptor II (QIAGEN, 20 sec on/off). The homogenate was filtered through a 40-μm cell strainer and centrifuged at 12,000 rpm for 20 minutes at 4 °C. NaCl was added to the supernatant to get a final concentration of 1.0 M NaCl. The supernatant was filtered through a 0.22 μm sterile filter and used for chitin affinity chromatography.

The stomach extract was purified on a 1.5 cm diameter, 10 mL column packed with chitin resin slurry (New England Biolabs). The column was pre-equilibrated with phosphate buffer (20 mM sodium phosphate buffer pH 7.0, 1.0 M NaCl) before the stomach extract was applied to the column. After washing with phosphate buffer (20 mM sodium phosphate buffer pH 7.0, 1.0 M NaCl), the chitinases were eluted with 100 mM acetic acid. A flow rate of 1 mL/min was used at all steps. Concentration and buffer exchange of the eluted chitinases to phosphate buffer (20 mM sodium phosphate buffer pH 7.0, 150 mM NaCl) was done using a 10 kDa centrifugal filter (Macrosep Advance Centrifugal Device, 10 kDa cutoff, Pall corporation). The purity of the eluted chitinases was examined with SDS-PAGE and Coomassie Brilliant Blue R-250 staining, and the proteins in the appearing bands were further identified by LC-MS at the local Proteomics Core Facility (NMBU). The protein concentration was determined with Quick Start Bradford Protein Assay (Bio-Rad) using Bovine Serum Albumin as standard.

### Confirmation of chitinase proteins with mass spectrometry

The protein bands in the de-stained gel were cut out using a clean scalpel blade. After trimming away unstained gel, the bands were further divided into 1-2 mm cubes and transferred to clean 0.2 mL PCR tubes. 100 μl of 50 % acetonitrile (ACN), 50 mM ammonium bicarbonate (ABC) was added to each tube, which was then incubated at room temperature with shaking for 10 minutes. After brief centrifugation, the liquid was aspirated and replaced with 200 μl 100 % ACN. The tubes were incubated at room temperature for 15 minutes and the liquid was removed by aspiration.

The in-gel reduction was performed by adding 50 μl 10 mM DTT, 50 mM ABC to the dried gel pieces, and incubating for 30 minutes at 56 °C in a thermocycler. Alkylation was performed by replacing the solution with 50 μl of 50 mM iodoacetamide, 50 mM ABC, and incubating in the dark for 20 min at room temperature.

After having removed the alkylation solution, 200 μl of 100% ACN was added and the tubes were incubated at room temperature for 15 minutes, followed by liquid removal and brief air drying of the gel pieces. The tubes were put on ice, and 30 μl ice-cold trypsin solution (13 ng/μl, in 50 mM ABC) was added to each tube. The gel pieces were allowed to swell for a total of 90 minutes on ice, with occasional checks to ensure that they were completely covered with the digestion solution. Finally, the tubes were transferred to a thermocycler and incubated overnight at 37 °C.

Trypsin digestion was terminated by adding 50 μl TFA solution (final concentration 0.2%), and the tubes were sonicated for 10 min in a water bath sonicator. After brief centrifugation, the liquid was transferred to a clean tube. 50 μl of 0.1% TFA was added to the gel pieces and the sonication step was repeated. After combining the two extracts, peptides were purified using STAGE spin-tips, as previously described ^54^. Eluted peptides were dried in an Eppendorf Concentrator Plus vacuum centrifuge and dissolved in loading solution (0.05% TFA, 2% ACN in Milli-Q water) before LC-MS/MS analysis.

Samples were loaded onto a trap column (Acclaim PepMap100, C_18_, 5 μm, 100 Å, 300 μm i.d. x 5 mm, Thermo Scientific) and backflushed onto a 50 cm analytical column (Acclaim PepMap RSLC C_18_, 2 μm, 100 Å, 75 μm i.d., Thermo Scientific). Starting conditions were 96 % solution A [0.1 % (v/v) formic acid], 4% solution B [80 % (v/v) ACN, 0.1 % (v/v) formic acid]. Peptides were eluted using a flow rate of 300 nl/min using a 70 min method, with the following gradient: from 3.2 to 10 % B in 3 minutes, 10 to 35 % B in 44 minutes and 35 to 60% B in 3 minutes, followed by a 5 min wash at 80 % B and a 15 min equilibration at 4% B. The Q-Exactive mass spectrometer was operated in data-dependent acquisition (DDA) mode using a Top10 DDA method, where acquisition alternates between orbitrap-MS and higher-energy collisional dissociation (HCD) orbitrap-MS/MS acquisition of the 10 most intense precursor ions. Only charge states 2-5 were selected for fragmentation, and the normalized collision energy (NCE) was set to 28. The selected precursor ions were excluded for repeated fragmentation for 20 seconds. The resolution was set to R=70,000 and R=35,000 for MS and MS/MS, respectively. Automatic gain control values were set to 3×10^6^ and 5×10^4^ for MS and MSMS, respectively, with a maximum injection time of 100 and 128 ms. The MS Raw files were analyzed using MaxQuant as described in the previous section.

### Chitinase assays with fluorogenic substrates

Chitinolytic activity was determined using a Chitinase Assay Kit (Fluorimetric, Sigma-Aldrich) with 4-Methylumbelliferyl N,N’-diacetyl- β -D-chitobioside (4-MU-GlcNAc_2_) and 4-Methylumbelliferyl β-D- N,N’,N’’-triacetylchitotriose (4-MU-GlcNAc_3_) according to the manufacturer’s protocol. To determine the activity in different salt concentrations 31 nM of purified chitinase was incubated with 100 μM 4-MU-GlcNAc_2_ in McIlvaine’s buffer (0.1 M citric acid and 0.2 M Na_2_HPO_4_, pH 5) with different concentrations of NaCl (0 – 0.6 M) in a volume of 100 μL at 37 °C for 15 minutes. To determine pH optimum, 31 nM (for assay with 4-MU-GlcNAc_2_) or 7.8 nM (for assay with 4-MU-GlcNAc_3_) of purified chitinase was incubated with 100 μM substrate in 0.1 M Gly-HCL buffer (pH 1.0 – 3.0) or McIlvaine’s buffer (0.1 M citric acid and 0.2 M Na_2_HPO_4_, pH 4.0 – 8.0) in a volume of 100 μL at 37 °C for 15 minutes. The reaction was stopped by the addition of 200 μL 0.4 M sodium carbonate. The fluorescence of the released 4-Methylumbelliferone (4-MU) was measured using a fluorimeter with excitation at 360 nm and emission at 450 nm and a 4-MU standard curve was used to quantify 4-MU resulting from the hydrolytic reaction. The measured fluorescence was corrected for hydrolysis of the substrate without the addition of an enzyme (blank). Each reaction was performed in triplicate.

### pH and stability assay

Purified chitinase enzyme or phosphate buffer (blank) was mixed with β-chitin in appropriate volumes of the following buffer solutions: 20 mM glycine-HCL (pH 2 and 3), 20 mM sodium acetate (pH 4 and 5), 20 mM sodium phosphate (pH 6), to yield final concentrations of 0.2 μM (chitinase) and 10 mg/mL (β-chitin). The reactions mixtures were incubated at 14 °C in an Eppendorf thermomixer at 1000 rpm and samples were taken after 0.5, 1, 3, 6 and 24 hours of incubation and filtered to remove β-chitin particles and thereby stop the reaction (0.45 μm vacuum filter, Merck Millipore). To adjust the samples for chromatography and to inactivate enzymes, H_2_SO_4_ was added to a final concentration of 20 mM in the filtrate. All reactions were run in triplicates. The end products were analyzed by HPLC (see below).

### Chitinase assay with α-chitin, shrimp shells, crab shells and black soldier fly pupae skin

Purified chitinase from Atlantic salmon (0.5 μM), purified ChiB from *Serratia marscecens* (0.5 μM) or phosphate buffer (blank) was mixed with α -chitin (10 mg/mL), shrimp shell (10 mg/mL), crab shell (10 mg/mL) and black soldier fly pupae skin (25 mg/mL) in 0.1 M sodium acetate (pH 4.8), and incubated at 14 °C in an Eppendorf thermomixer at 1000 rpm (all concentrations noted in parenthesis indicate final concentrations in the reaction mixtures). Samples were withdrawn after 6 and 24 hours of incubation, filtered by a 0.45 μm filter to remove the substrate particles from the reaction mixture and thereby stop the reaction. The enzyme activity was inactivated by addition H_2_SO_4_ to a final concentration of 20 mM in the filtrate. All reactions were run in triplicates. The end products were analyzed by HPLC (see below).

### High-performance liquid chromatography (HPLC)

Concentrations of mono- and disaccharides of *N*-acetylglucosamine (GlcNAc and GlcNAc_2_) were determined as previously described ^55^.

### Availability of data

The proteomics data have been deposited to the ProteomeXchange consortium (http://proteomecentral.proteomexchange.org) via the PRIDE ^56^ partner repository with the dataset identifier PXD030291.

