## Supplementary Table 1 for "The gastric mucosa of Atlantic salmon (*Salmo salar*) is abundant in highly active chitinases"

**Supplementary Table 1**. Top 20 proteins with putative secretion encoding signal peptides in Atlantic salmon stomach mucosa, identified with label-free quantitative proteomics.

| LFQ intensity | SE | Uniprot ID | Protein name |
| --- | --- | --- | --- |
| 33.26 | 0.84 | A0A1S3L8D8 | **Chitinase (Chia.3)** |
| 32.30 | 0.55 | A0A1S3NDX6 | Pepsin- A-like |
| 31.57 | 0.53 | A0A1S3MFN1 | **Chitinase (Chia.7)** |
| 30.80 | 0.72 | A0A1S3L8T9 | **Chitinase (Chia.4)** |
| 29.95 | 0.04 | A0A1S3MIE3 | Pepsin- A-like |
| 29.85 | 1.69 | B5X6A6 | Cystatin |
| 29.12 | 0.92 | B9ENV9 | Fish-egg lectin |
| 29.09 | 2.21 | A0A1S3PXG0 | IgGFc-binding protein-like |
| 28.65 | 0.16 | A0A1S3R1S1 | Serotransferrin |
| 28.33 | 0.65 | A0A1S3PQV6 | Hemopexin |
| 26.95 | 0.24 | B5XBH3 | Apolipoprotein |
| 26.58 | 0.34 | A0A1S3KR74 | Metalloendopeptidase |
| 25.88 | 0.48 | A0A1S3MUF1 | Uncharacterized |
| 25.60 | 1.07 | A0A1S3LBJ3 | Uncharacterized |
| 24.97 | 0.12 | B5XB02 | Chymotrypsin B |
| 24.78 | 0.31 | Q2V6Q7 | Anterior gradient 2-like |
| 24.39 | 0.46 | A0A1S3KS25 | Endoplasmin-like |
| 24.17 | 0.06 | A0A1S3S2K4 | Pancreatic secretory granule membrane major glycoprotein-like |
| 23.93 | 0.98 | B5X8N0 | Carboxypeptidase A1 |
| 23.06 | 0.30 | A0A1S3KK24 | Histidine-rich glycoprotein-like |
